# Associations of waking cortisol with DHEA and testosterone across the pubertal transition: Effects of threat-related early life stress

**DOI:** 10.1101/691279

**Authors:** Lucy S. King, Madelaine G. Graber, Natalie L. Colich, Ian H. Gotlib

**Affiliations:** Stanford University; University of Washington

**Keywords:** cortisol, DHEA, testosterone, puberty, adolescence, early life stress

## Abstract

Atypical regulation of the hypothalamic-pituitary-adrenal (HPA) axis is a putative mechanism underlying the association between exposure to early life stress (ELS) and the subsequent development of mental and physical health difficulties. Recent research indicates that puberty is a period of HPA-axis plasticity during which the effects of exposure to ELS on cortisol regulation may change. In particular, increases in the sex hormones that drive pubertal maturation, including dehydroepiandrosterone (DHEA) and testosterone, may be implicated in pubertal recalibration of cortisol regulation. In the current study, we examined the associations among levels of objectively-rated threat-related ELS and salivary waking cortisol, DHEA, and testosterone in a sample of 178 adolescents (55% female) who were in early puberty at baseline (Tanner stages 1-3; mean Tanner stage[SD]=1.93[0.64]; mean age[SD]=11.42[1.04]) and were followed up approximately two years later (mean Tanner stage[SD]=3.46[0.86]; mean age[SD]=13.38[1.06]). Using multi-level modeling, we disaggregated the effects of between-individual levels and within-individual increases in pubertal stage and sex hormones on change in cortisol. Controlling for between-individual differences in average pubertal stage, the association between levels of cortisol and DHEA was more strongly positive among adolescents who evidenced greater within-individual increases in pubertal stage across time. Both higher average levels and greater within-individual increases in DHEA and testosterone were associated with increases in cortisol across time, indicating positive coupling of developmental changes in these hormones; however, coupling was attenuated in adolescents who were exposed to more severe threat-related ELS prior to puberty. These findings advance our understanding of the development of the HPA-axis and its association with childhood environmental risk during puberty.

## 1. Introduction

The hypothalamic-pituitary-adrenal (HPA) axis mediates physiological responses to stress (Gunnar and Quevedo, 2007), and functioning of the HPA axis has pervasive effects on mental and physical health (Koss and Gunnar, 2018). A burgeoning literature implicates atypical cortisol regulation, both diurnally and in response to stress, in the development of psychopathology (e.g., Adam et al., 2017; Colich, Kircanski, Foland-Ross, & Gotlib, 2015; Essex et al., 2011). Further, many studies have linked exposure to early life stress (ELS; with “early” defined as occurring during childhood) with subsequent cortisol dysregulation (e.g., Bernard, Frost, Bennett, & Lindhiem, 2017; Bunea, Szentágotai-Tătar, & Miu, 2017).

The associations among cortisol, ELS, and health are complex. For example, the effects of cortisol on health follow an inverted U-shaped curve (Sapolsky, 1997). Because moderate levels of cortisol are critical for adaptive responses to stress, atypically low or “blunted” levels of cortisol may increase risk for poor health; in contrast, chronically high levels of cortisol may indicate the loss of neurobiological resilience (McEwen, 2019). In addition, the effects of ELS on cortisol regulation may depend on developmental stage. Specifically, there is increasing evidence that puberty is a period of HPA-axis plasticity, and that the effects of ELS on cortisol regulation change with pubertal maturation. In two cross-sectional studies, ELS in the form of institutionalization prior to international adoption was associated with blunted cortisol responses to awakening (Quevedo et al., 2012) and to social stress (DePasquale et al., 2019) in children in *early* puberty. In these studies, however, the cortisol responses of previously institutionalized children in *later* puberty were indistinguishable from those of never-adopted children (DePasquale et al., 2019; Quevedo et al., 2012). In another cross-sectional study, we found that the association between ELS and cortisol responses to awakening in adolescents did not dissipate but *differed* in participants as a function of pubertal maturation. Specifically, greater severity of exposure to a variety of life stressors prior to puberty (i.e., parental divorce, maltreatment, community violence) was associated with a blunted cortisol awakening response in early puberty (King et al., 2017) but a heightened cortisol response in later puberty (King et al., 2017). Recently, using an accelerated longitudinal design, Gunnar et al. (2019) found that within-individual increases in pubertal stage moderated the association between a history of institutionalization and cortisol responses to social stress; the cortisol responses of previously institutionalized children became more similar to those of never-adopted children as they progressed through puberty. The biological mechanisms underlying changes in the effects of ELS on cortisol regulation during puberty remain unclear (DePasquale et al., 2019; Romeo, 2013).

Researchers have long known that puberty is characterized by dramatic increases in sex hormones that are responsible for the development of secondary sex characteristics and the attainment of sexual maturation. It is possible that changes in cortisol regulation and its association with ELS during puberty are driven by these increases in sex hormones. Adrenarche is the first major sign of puberty, involving rises in adrenal hormones, including dehydroepiandrosterone (DHEA), beginning at age 6-8 years in girls and approximately one year later in boys (Dorn and Biro, 2011). Increases in DHEA contribute in part to gonadarche, or the re-activation of the hypothalamic-pituitary-gonadal (HPG) axis and attendant rises in testosterone and estradiol (Dorn and Biro, 2011; Marceau et al., 2015a). The synthesis and metabolism of cortisol is related to that of DHEA and testosterone. Both cortisol and DHEA are produced by the HPA axis and testosterone is produced predominantly by the HPG axis (Marceau, Ruttle, Shirtcliff, Essex, et al., 2015a; although a proportion of tesosterone is produced by the adrenal glands in both sexes and this proportion is higher in girls [Bordini & Rosenfield, 2011]). Cortisol, DHEA, and testosterone follow a similar diurnal pattern of higher morning levels that decline throughout the course of the day. Further, all three hormones show increases in response to acute stress (Hucklebridge et al., 2005; Marceau et al., 2014; Matchock et al., 2007). The covariation of cortisol and DHEA may be protective. Although the neurobiological actions of DHEA are complex and higher concentrations of DHEA are not necessarily beneficial for health, DHEA has anti-glucocorticoid effects (Maninger et al., 2009). Specifically, DHEA and cortisol appear to have opposing regulatory functions such that DHEA protects against the neurotoxic effects of elevated cortisol (Kamin and Kertes, 2016). In adults, the activation of the HPA axis appears to suppress the HPG axis (Mastorakos et al., 2006); however, emerging research suggests that this is not the case in early adolescence (Marceau et al., 2014).

A small number of studies have examined the associations of sex hormones with cortisol during adolescence. These studies disaggregate within- versus between-individual effects of sex hormones on cortisol, or the effects of changes in sex hormones on changes in cortisol within an individual (hereafter referred to as “coupling”) versus the effects of average levels of sex hormones on levels of cortisol between individuals (Marceau et al., 2015b). DHEA and testosterone evidence positive diurnal coupling with cortisol, such that when adolescents display higher levels of cortisol they also display higher levels of DHEA and testosterone (Black et al., 2018; Dismukes et al., 2015b; Johnson et al., 2014; Marceau et al., 2015b). Marceau et al. (2014) found that cortisol was also positively coupled with DHEA and testosterone in response to stress; however, the strength of coupling varied based on age and sex. Specifically, cortisol– *DHEA* coupling was stronger in older boys, whereas cortisol–testosterone coupling was weaker in older boys. Only one previous study has investigated the association of cortisol with sex hormones across development. Ruttle, Shirtcliff, Armstrong, Klein, & Essex (2013) assessed waking levels of cortisol, DHEA, and testosterone on three days from early to middle adolescence, finding that cortisol–DHEA coupling across the three days was more strongly positive in later adolescence, whereas cortisol–testosterone coupling was more negative in later adolescence. No previous study has investigated how changes in pubertal maturation influence the associations between levels of sex hormones and cortisol or whether developmental increases in sex hormones during puberty are coupled with changes in cortisol during this period.

Previous studies of the association of cortisol with DHEA and testosterone have recruited adolescents on the basis of age rather than of pubertal stage, limiting the ability to examine the relation between pubertal maturation—as defined by both external physical and hormonal changes—and changes in cortisol. Boys and girls differ significantly in pubertal timing, with girls typically experiencing the onset of puberty 1.5 years earlier than do boys (Negriff and Susman, 2011). Given these sex differences in pubertal timing, age-matched samples of adolescent boys and girls are almost certain to be confounded by sex differences in pubertal stage. This limitation makes it difficult to isolate the dynamics of cortisol and sex hormones that are specific to the pubertal transition in both boys and girls. Consequently, estimates of the effects of environmental risk on hormonal coupling are similarly limited. Further, because individuals enter puberty at different ages, recruitment based on age leads to wide variation in pubertal stage at the baseline assessment. Some young adolescents may already be in the later stages of puberty or be fully developed at baseline. For these adolescents, follow up assessments do not capture the pubertal transition.

### 1.1 The Current Study

In the current study we used a longitudinal design with two timepoints to disaggregate the effects of between-individual levels and within-individual increases in pubertal stage and sex hormones (DHEA and testosterone) on cortisol across the transition from early (Time 1 [T1]; mean age=11.42; mean Tanner stage=1.92 [range 1-3]) to later puberty (Time 2 [T2]; ∼2 years later; mean age=13.38; mean Tanner stage=3.44 [range 1-5]). In contrast to previous studies, we recruited boys and girls on the basis of early pubertal stage (self-reported Tanner stage ≤ 3) rather than age. All boys and girls were in early puberty at the initial timepoint and were followed up approximately two years later, capturing a period of development during which pubertal maturation occurs. In addition, we evaluated the effect of exposure to ELS on the effect of within-person developmental increases in sex hormones on changes in cortisol across time (i.e., the coupling of developmental changes). The two previous cross-sectional studies of the coupling of sex hormones and cortisol in adolescence in the context of childhood adversity focused on ELS in the form of childhood maltreatment (Dismukes et al., 2015b; Simmons et al., 2015). In a cross-sectional study of nine-year-old children, Black et al. (2017) operationalized ELS as a count of stressful life events, including moving, financial problems, and abuse. The single previous longitudinal investigation operationalized ELS as a composite of parent-reported depression, perceived stress, and family expressed anger (Ruttle et al., 2015). There is evidence that different forms of ELS have differential effects on psychobiological development (King et al., 2019; Lambert et al., 2017; Sheridan et al., 2017) and, in particular, that exposure to environmental threat (as opposed to deprivation) accelerates biological development (Colich et al., 2019; Del Giudice et al., 2011). Therefore, measures that combine exposure to different forms of ELS make it difficult to identify directional hypotheses or to determine what form of stress confers risk. In the current study, we focused on threat-related ELS (i.e., exposure to domestic and community violence) as opposed to other forms of adversity in order to increase the specificity of our hypotheses.

We had three objectives in this study. First, we examined the effects of between-individual differences in average pubertal stage and within-individual increases in pubertal stage across timepoints on the association of waking levels of cortisol with waking levels of DHEA and testosterone. Based on findings from cross-sectional studies that cortisol–DHEA coupling is positive during adolescence (Dismukes et al., 2015b; Johnson et al., 2014; Marceau et al., 2015b), and the results of a longitudinal study indicating that cortisol–DHEA coupling is more strongly positive in later adolescence (Ruttle et al., 2015), we hypothesized that waking levels of cortisol and DHEA are positively associated, and that this positive association is *stronger* in adolescents who are, on average, further along in pubertal development (later puberty) and who evidence greater increases in pubertal stage across timepoints. In contrast, based on findings that cortisol–testosterone diurnal coupling is positive during adolescence (Dismukes et al., 2015b; Johnson et al., 2014; Marceau et al., 2015b), but negative in later adolescence (Ruttle et al., 2015), we hypothesized that waking levels of cortisol and testosterone are positively associated, but that this association is *weaker* in adolescents who are, on average, further along in pubertal development (later puberty), and who evidence greater increases in pubertal stage across timepoints.

Second, we examined the effects of within-individual changes in waking DHEA and testosterone on change in waking levels of cortisol across timepoints (i.e., coupling of developmental increases) when controlling for between-individual levels of DHEA and testosterone. DHEA and testosterone increase with pubertal maturation (Dorn and Biro, 2011); further, cortisol and DHEA have been found to be positively coupled throughout adolescence whereas cortisol and testosterone have been found to be more weakly or negatively coupled in later adolescence (Ruttle et al., 2015). Therefore, we hypothesized that greater increases in DHEA, but not in testosterone, across development are coupled with increases in cortisol across development.

Third, we examined the between-individual effect of ELS in the form of the severity of exposure to threat-related ELS (i.e., domestic and community violence) on the coupling of developmental increases in DHEA and testosterone with changes in cortisol across timepoints. Given evidence that exposure to environmental threat (as opposed to deprivation) accelerates biological development (Colich et al., 2019; Del Giudice et al., 2011), we hypothesized that greater severity of threat-related ELS is associated with stronger positive coupling between change in cortisol and change in DHEA, and with weaker positive coupling between change in cortisol and change in testosterone across development.

## 2. Methods

### 2.1 Participants

Participants were 214 early adolescents and their parents who were recruited from the community to participate in a longitudinal study of the psychobiological effects of ELS across the transition from early puberty (T1) to later puberty (T2) (Humphreys, Kircanski, Colich, & Gotlib, 2016; King, Humphreys, Camacho, & Gotlib, 2018). In this study, *early puberty* was defined by self-reported Tanner stages 1-3. Participants were recruited from the geographic area surrounding Stanford University through local and media postings. Inclusion criteria at T1 were that the adolescents were between 9 and 13 years of age and proficient in spoken English. Exclusion criteria at T1 included a history of major neurological or medical illnesses, severe learning disabilities that would affect comprehension of study procedures, and, for females, the onset of menses. In addition, participants were selected based on their eligibility to participate in a magnetic resonance imaging (MRI) scan (no metal implants). Given the focus of the study on pubertal development, boys and girls were matched on self-reported pubertal stage using Tanner staging (see “Pubertal Stage,” below). To be included in the current analyses, we required that participants be in early puberty at T1 (Tanner stages 1-3), resulting in the exclusion of 14 participants. We excluded an additional 24 participants who did not provide a waking saliva sample that yielded values for cortisol, DHEA, or testosterone at either T1 or T2. Therefore, the final sample for the current analyses was 178 adolescents (55% female), 171 of whom provided data at T1 and 147 of whom provided data at T2. Descriptive statistics for the final study sample are presented in Table 1. There were no significant differences at T1 between participants who were and who were not included in the final sample with respect to threat-related ELS (described below), family income-to-needs ratio, pubertal stage, or body mass index (BMI). The sample for this study overlaps with that of an earlier study in which diurnal cortisol was examined only at the T1 assessment (King et al., 2017); the cortisol and sex hormone data reported in this study were not reported in the earlier study.

**Table 1.** Descriptive statistics of the study sample.

| Measure | Sex | N |  | Mean (SD) |  |
| --- | --- | --- | --- | --- | --- |
|  |  | T1 | T2 | T1 | T2 |
| Age | Female | 94 | 76 | 11.07 (0.95) | 12.96 (0.96) |
|  | Male | 77 | 71 | 11.87 (0.97) | 13.82 (0.96) |
| Pubertal Stage | Female | 94 | 75 | 2.00 (0.68) | 3.42 (0.79) |
|  | Male | 77 | 63 | 1.85 (0.59) | 3.48 (0.92) |
| BMI | Female | 92 | 62 | 18.31 (4.04) | 20.15 (4.86) |
|  | Male | 74 | 56 | 19.04 (3.38) | 20.78 (3.94) |
| Cortisol $\mu\text{g/dL}$ | Female | 81 | 72 | 0.28 (0.18) | 0.26 (0.16) |
|  | Male | 70 | 68 | 0.20 (0.11) | 0.29 (0.13) |
| DHEA pg/mL | Female | 94 | 74 | 155.81 (128.12) | 168.31 (130.36) |
|  | Male | 76 | 70 | 187.24 (330.35) | 181.31 (118.90) |
| Testosterone pg/mL | Female | 93 | 76 | 53.03 (20.75) | 61.60 (28.73) |
|  | Male | 75 | 71 | 61.90 (33.77) | 134.41 (76.75) |
| Saliva collection time | Female | 85 | 71 | 7.30 (1.20) | 8.05 (1.67) |
|  | Male | 67 | 65 | 7.64 (1.29) | 8.18 (1.71) |
| Threat-related ELS | Female | 94 | 73 | 2.03 (2.13) |  |
|  | Male | 77 | 67 | 1.97 (1.98) |  |
| Income-to-needs ratio | Female | 85 | 67 | 1.32 (0.53) |  |
|  | Male | 71 | 60 | 1.33 (0.52) |  |
| Race/ethnicity |  |  |  |  |  |
| White | Female | 46 |  |  |  |
|  | Male | 35 |  |  |  |
| Biracial | Female | 16 |  |  |  |
|  | Male | 20 |  |  |  |
| Asian or Asian American | Female | 15 |  |  |  |
|  | Male | 7 |  |  |  |
| Hispanic | Female | 8 |  |  |  |
|  | Male | 3 |  |  |  |
| Black or African American | Female | 5 |  |  |  |
|  | Male | 5 |  |  |  |
| Other | Female | 4 |  |  |  |
|  | Male | 6 |  |  |  |
| Not reported | Female | 0 |  |  |  |
|  | Male | 1 |  |  |  |

**Notes.** T1 = baseline assessment in early puberty. T2 = follow-up assessment approximately 2 years later BMI = body mass index. ELS = early life stress. Although raw hormone values are presented in Table 1, hormone values were natural log-transformed to reduce positive skew for the data analyses. Saliva collection time is in hours from midnight.

### 2.2 Procedure

The Stanford University Institutional Review Board approved the protocol for this study. In an initial telephone call, research staff provided information about the study to families and screened participants for inclusion/exclusion criteria. Eligible families were then invited to attend a laboratory session during which staff obtained consent from parents and assent from adolescents. In this session, adolescents reported their pubertal stages, and both parents and adolescents completed interview and questionnaire measures about the child and family. At the end of the session, staff provided families with kits and instructions to collect saliva samples at home for the assessment of waking hormone levels. Families returned the samples to the laboratory at a subsequent visit. These procedures were repeated at a T2 laboratory session that occurred an average of 2 years later (mean[SD]=1.96[0.31] years; range: 1.18-2.88). The timing of the T2 assessment varied for a variety of reasons (reviewed in the Supplementary Material) related to the challenges of collecting longitudinal data in a large adolescent sample.

### 2.3 Measures

#### 2.3.1 Pubertal stage

In order to match boys and girls based on pubertal stage at T1, we measured pubertal development using self-report Tanner staging (Marshall and Tanner, 1968; Morris and Udry, 1980). Self-report Tanner staging scores are moderately correlated with physicians’ physical examinations of pubertal development (Dorn and Biro, 2011). Participants reported their pubertal stage by selecting how closely their pubic hair and breast/testes resembled an array of schematic drawings on a scale of 1 (prepubertal) to 5 (postpubertal). For the purposes of this study, we used the average of the pubic hair and breast/testes Tanner scores to index overall pubertal stage. Average Tanner scores ranged from 1-3 at T1 and from 1-5 at T2. Five children did not report increases in Tanner stage from T1 to T2 (mean increase[SD]=1.50[0.77]). Distributions of Tanner scores at each timepoint and change in Tanner scores from T1 to T2 are presented in the Supplementary Material. Nine participants did not complete the Tanner staging questionnaire at T2.

#### 2.3.2 Severity of threat-related stress

As previously described (King et al., 2019, 2017), at T1, participants were interviewed about their lifetime exposure to 30 types of stressors using a modified version of the Traumatic Events Screening Inventory for Children (Ribbe, 1996). A panel of three coders, blind to the adolescents’ reactions and behaviors during the interview, then rated the objective severity of each type of stressor endorsed on a scale of (0 = non-event or no impact; 4 = extremely severe impact; ICC = 0.99). To quantify the severity of threat-related ELS, we summed the maximum objective severity ratings for the events listed in in Table 2 (see https://github.com/lucysking/els_stress_interview for scoring script). We selected those events that were interpersonal in nature and consistent with Sheridan and McLaughlin’s (2014) definition of threat as “the presence of an atypical (i.e., unexpected) experience characterized by actual or threatened death, injury, sexual violation, or other harm to one’s physical integrity” (p. 580). Adolescents were interviewed again at the T2 assessment about exposure to the same 30 types of stressors since the T1 assessment (see Supplementary Material for descriptive statistics).

**Table 2.** Endorsement of lifetime threat-related stressors occurring up to early puberty (T1)

| Type of ELS | N<br>endorsed | %<br>endorsed | Mean (SD)<br>Severity |
| --- | --- | --- | --- |
| Family verbal conflict | 72 | 41 | 1.96(0.72) |
| Bullying | 60 | 34 | 1.35(0.29) |
| Community violence | 21 | 12 | 0.86(0.64) |
| Community instability | 17 | 10 | 0.97(0.60) |
| Domestic violence | 12 | 7 | 2.39(0.84) |
| Community verbal conflict | 9 | 5 | 0.78(0.51) |
| Emotional abuse | 9 | 5 | 1.50(0.87) |
| Physical abuse | 8 | 5 | 2.19(1.00) |
| Mugging or robbery | 5 | 3 | 1.10 (0.82) |
| War or terrorism | 5 | 3 | 0.50 (0.00) |
| Sexual abuse | 3 | 2 | 3.00(1.73) |
| Threats of domestic violence | 3 | 2 | 1.67(1.04) |
| Kidnapping | 2 | 1 | 1.00(0.00) |
| Threats of physical abuse | 2 | 1 | 2.00(0.00) |
| Witness sexual abuse | 1 | 1 | 2.00(NA) |

**Notes.** Stressors coded as "community instability" included community-level threats (e.g., bomb/active shooter threats at school, hearing gun shots in neighborhood). Stressors coded as "war or terrorism" included witnessing events live on television.

#### 2.3.3 Waking cortisol, DHEA, and testosterone

Salivary hormonal assays were conducted for cortisol, DHEA and testosterone. At each timepoint, participants were asked to provide a saliva sample through passive drool immediately upon awakening (prior to eating, drinking, or brushing teeth). Participants recorded collection time and placed the saliva samples in their home freezer after collection. Contemporaneous with the saliva samples, participants reported their use of medications, including corticosteroids. After participants returned the samples to the laboratory, samples were transferred to a −20°C freezer and subsequently shipped on dry ice to Salimetrics, LLC (State College, PA), where they were assayed for cortisol, DHEA, and testosterone using a high sensitivity enzyme immunoassays. The average intra- and inter-assay coefficients of variation for cortisol were 4.60% and 6.00%, respectively. The average intra- and inter-assay coefficients variation for DHEA were 5.55% and 8.20%, respectively. The average intra- and inter-assay coefficients variation for testosterone were 4.60% and 9.85%, respectively. Additional information concerning the assays is provided in the Supplementary Material. We winsorized cortisol, DHEA, and testosterone values that were +/− 3SD from the mean (within timepoint and sex), and then log-transformed the values at each timepoint to correct for positive skew. To quantify reliability of the hormone values, we calculated the intra-class correlation coefficients (ICCs; two-way mixed effects) across T1 and T2. Although we anticipated developmental change in hormones between T1 and T2, values evidenced moderate rank-order stability within individuals and agreement across timepoints: ICC_CORTISOL_=.33, ICC_DHEA_=.69, ICC_TESTOSTERONE_=.58.

### 2.4 Data analysis

All analyses were conducted in R (R Core Team, 2018). We used multi-level modeling (also known as mixed-effects or hierarchical linear modeling; Woltman, Feldstain, Mackay, & Rocchi, 2012) to test our hypotheses. We implemented these models using the function “lmer” in the “lme4” package, and we used the package “lmerTest” to calculate degrees of freedom and p-values (Bates et al., 2014; Kuznetsova et al., 2016). Data and analysis scripts are available at: https://github.com/lucysking/els_cort_dhea. All variables were standardized prior to analysis such that estimates reflect the predicted change in the dependent variable for a 1-SD change in the independent variable.

#### 2.4.1 Objective 1: Examine within- and between-individual effects of pubertal stage on the associations of levels of cortisol with levels of sex hormones

To distinguish within- and between-individual effects of pubertal stage on the associations of waking levels of cortisol with waking levels of DHEA and testosterone, we calculated two terms: 1) longitudinal change (Δ) in pubertal stage from T1 to T2, modeled by centering pubertal stage around each adolescent’s mean across T1 and T2 (i.e., person-mean-centering); and 2) average pubertal stage compared to other adolescents, modeled by averaging each adolescent’s pubertal stage across timepoints (Curran and Bauer, 2011; see Gunnar et al., 2019 for a similar approach). This model was fit as follows, controlling for the random effect of participant intercepts:

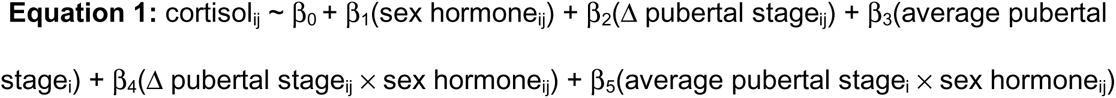

In this context, a significant interaction between Δ pubertal stage and the sex hormone (DHEA or testosterone) indicates that the magnitude of the association between the sex hormone and cortisol depends on the degree of pubertal maturation between T1 and T2, regardless of the average level of maturity. In contrast, a significant interaction between average pubertal stage and the sex hormone indicates that the magnitude of association between the sex hormone and cortisol depends on between-individual differences in average level of maturity across timepoints.

#### 2.4.2 Objective 2: Examine coupling of developmental increases in sex hormones with changes in cortisol

We next examined whether developmental increases in sex hormones between T1 and T2 are coupled with changes in cortisol across this period, controlling for between-individual differences in average levels of sex hormones across timepoints. Using a similar approach as in Objective 1 to disaggregate within- and between-individual effects. we calculated two terms: 1) longitudinal change (Δ) in each adolescent’s sex hormone (DHEA or testosterone), modeled by centering the sex hormone around each adolescent’s mean across T1 and T2; and 2) average level of the sex hormone compared to other adolescents, modeled by averaging each adolescent’s sex hormones across T1 and T2. This model was fit as follows, controlling for the random effect of participant intercepts:

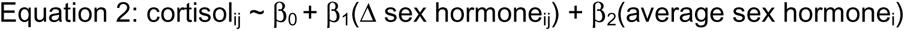

In this context, a significant effect of Δ sex hormone indicates that within-individual increases in the sex hormone are associated with changes in cortisol from T1 to T2, regardless of the average level of the sex hormone across timepoints.

#### 2.4.3 Objective 3: Examine the effect of the severity of threat-related ELS on the coupling of developmental increases in sex hormones with changes in cortisol

Finally, we tested the between-individual effects of the severity of threat-related ELS on the coupling of developmental changes in sex hormones and cortisol by adding interactions with the severity of threat-related ELS to the model above. Specifically, we fit the following model:

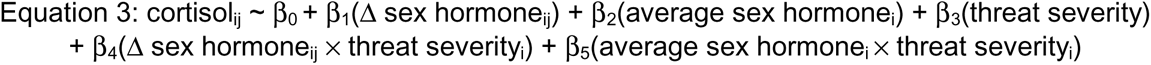

In this context, a significant interaction between the Δ sex hormone and the severity of threat-related ELS indicates that the coupling of within-individual changes in the sex hormone with changes in cortisol depends on between-individual differences in level of threat severity.

To examine the impact of potential covariates, including interval in years between T1 and T2, use of medication, sex, BMI, and age at T1 on the effects of interest, we first conducted formal model fitting (Chambers, 1992) in which we tested whether each of the covariates significantly improved the model fit using likelihood ratio tests. Next, we conducted sensitivity analyses including the covariates that improved model fit. In addition, we tested separate models in which we removed observations from participants who were taking corticosteroid medications at the time of hormone collection. Results were highly similar in these separate models. To determine whether the effects of stress occurring earlier in life (i.e., prior to T1) were distinct from those of stress occurring during the transition through puberty (i.e., events occurring between the T1 and T2 assessments reported at T2), we tested separate models in which we replaced threat-related ELS in Equation 3 by the severity of threatening experiences occurring between T1 and T2.

Finally, given sex differences in the production and function of cortisol, DHEA, and testosterone, we followed previous research investigating hormonal dynamics in adolescence (Simmons et al., 2015) by running the models specified in Equations 1-3 separately for boys and girls.

## 3. Results

### 3.1 Sample characteristics

Characteristics of the sample are presented in Table 1. At T1, 15% of the adolescents reported using medications contemporaneous with saliva samples; 4% reported using a corticosteroid. At T2, 20% of adolescents reported using medications; 6% reported using a corticosteroid. None of the girls in the study reported using hormonal birth control at either timepoint. Severity of threat-related ELS at T1 was not correlated significantly with adolescent age, pubertal stage, BMI, or years between the T1 and T2 assessments. DHEA was not correlated with pubertal stage at T1 (*r*(168)=.11, *p*=.144) but was positively correlated with pubertal stage at T2 (*r*(134)=.38, *p*<.001). Testosterone was weakly positively correlated with pubertal stage at T1 (*r*(166)=.19, *p*=.015) and more strongly positively correlated with pubertal stage at T2 (*r*(136)=.39, *p*<.001). Finally, cortisol was not correlated with pubertal stage at T1 (*r*(149)=.01, *p=*.904) but was weakly positively correlated with pubertal stage at T2 (*r*(130)*=*.20, *p*=.022). As expected based on the design of the study, self-reported pubertal stage increased significantly from T1 to T2 (*t*(134)=22.67, *p*<.001), as did BMI (*t*(109)=6.69, *p*<.001), and age (*t*(141)=1.98, *p*<.001).

Statistics for significant differences between boys and girls in the study measures are presented in the Supplementary Material. Boys and girls did not differ significantly in severity of threat-related ELS, in the length of the interval between T1 and T2, or in pubertal stage, BMI, or DHEA at either timepoint. Girls had significantly higher levels of cortisol at T1 than did boys. Further, multi-level models indicated that sex interacted with Δ pubertal stage to explain cortisol; within-individual increases in pubertal stage between T1 and T2 were associated with increases in cortisol in boys but not in girls. At T1, boys and girls did not differ significantly in levels of testosterone; however, at T2, boys had significantly higher levels of testosterone than did girls. Sex interacted with Δ pubertal stage to explain testosterone; within-individual increases in pubertal stage were associated with higher testosterone for both boys and girls, but this association was stronger in boys than in girls. There were no sex differences in levels of DHEA at either timepoint or in the within- and between-individual effects of pubertal stage on change in DHEA across timepoints. Instead, greater Δ pubertal stage was associated with larger increases in DHEA between T1 and T2 for both boys and girls. Sex differences in cortisol, DHEA, and testosterone at each timepoint and as a function of Δ pubertal stage are presented in Figure 1.

**Figure 1.**
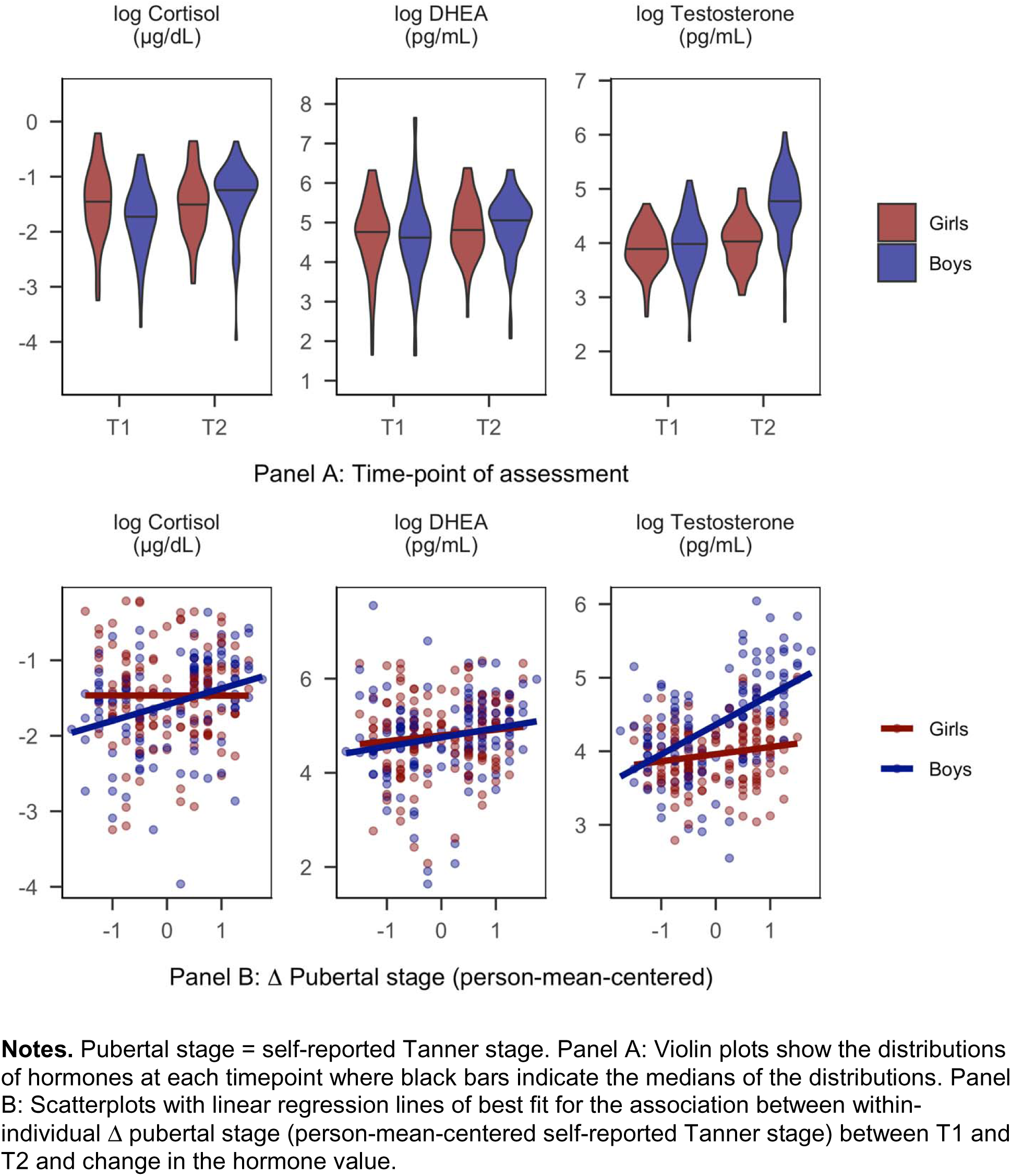
Sex differences in cortisol, DHEA, and testosterone.

### 3.2 Objective 1: Examine within- and between-individual effects of pubertal stage on the associations of levels of cortisol with levels of sex hormones

#### 3.2.1 DHEA

Waking levels of DHEA interacted with Δ pubertal stage to explain variation in waking levels of cortisol (β=0.15, SE=0.06, *t*(178.72)=2.36, *p*=.019, 95% CI[0.01, 0.26]), but did not interact with average pubertal stage (β=0.04, SE=0.07, *t*(236.88)=0.56, *p*=.577, 95% CI[−0.11, 0.19]). Supporting our hypothesis, simple effects analyses at +/− 1SD of mean Δ pubertal stage indicated that among adolescents with larger increases in pubertal stage between T1 and T2, DHEA was more strongly positively associated with cortisol (β=0.64, SE=0.09, *t*(235.73)=7.02, *p*<.001, 95% CI[0.44, 0.82]) than it was among adolescents with smaller increases in pubertal stage (β=0.39, SE=0.08, *t*(234.26)=5.22, *p*<.001, 95% CI[0.24, 0.55]). Results of formal model fitting indicated that the interaction between sex and Δ pubertal stage improved model fit. Results were highly similar when controlling for this interaction.

In a separate model conducted within boys only, the interaction between DHEA and Δ pubertal stage was larger in effect size than it was in the full model that included both boys and girls (β=0.25, SE=0.09, *p*=.009, 95% CI[0.07, 0.44]). In contrast, in a separate model conducted within girls only, the interaction between DHEA and Δ pubertal stage approached zero (β=0.08, SE=0.08, 95% CI[−0.09, 0.24]). The simple associations between DHEA and cortisol at T1 and T2 for each sex are presented in Figure 2, Panel A.

**Figure 2.**
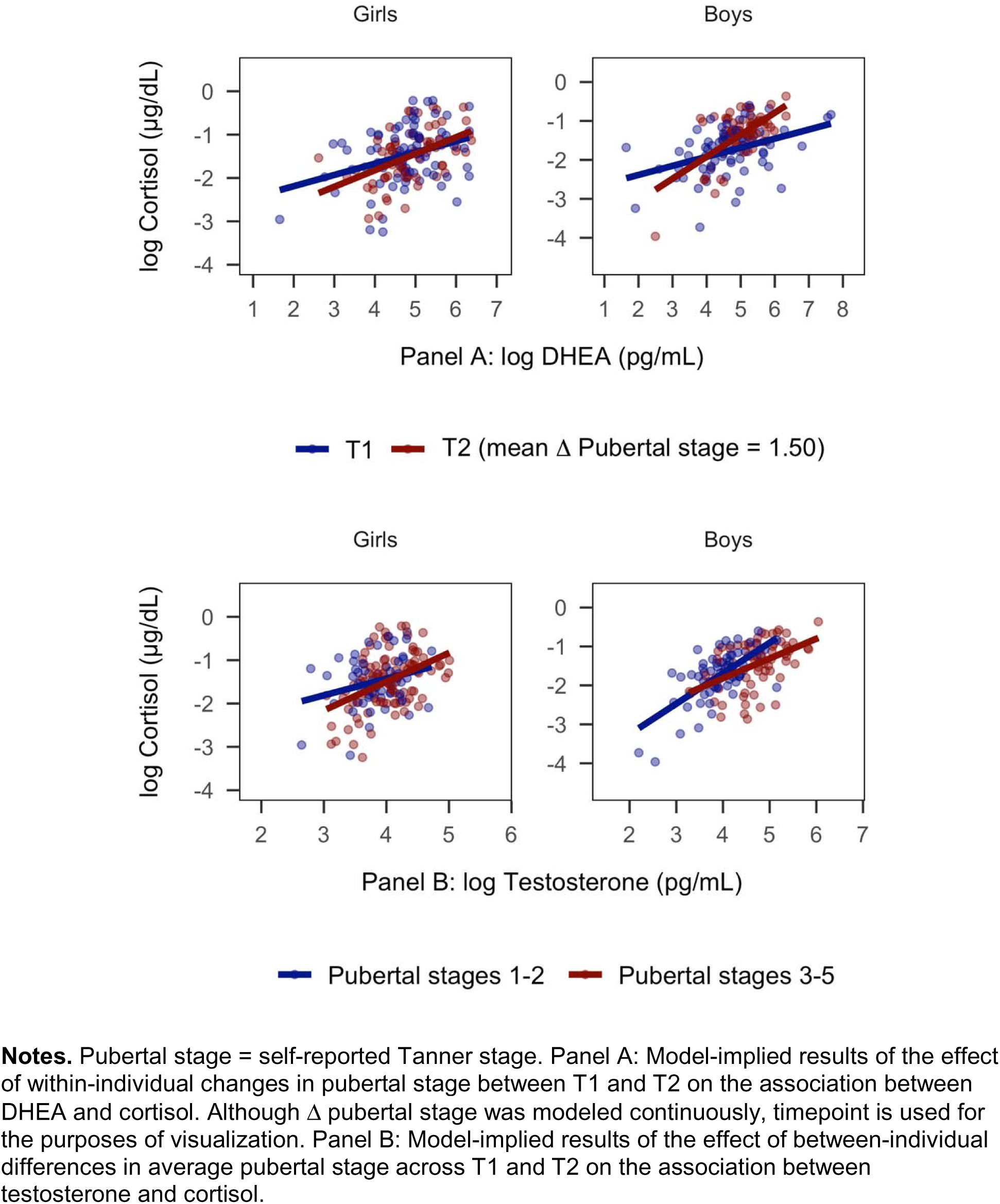
Within-individual increases in pubertal stage and between-individual differences in average maturity moderate the associations of cortisol with DHEA and testosterone.

#### 3.2.2 Testosterone

Waking levels of testosterone did not interact with Δ pubertal stage to explain variation in levels of cortisol (β=0.02, SE=0.06, *t*(193.08)=0.35, *p*=.728, 95% CI[−0.10, 0.14]). Instead, there was a significant interaction between testosterone and average pubertal stage (β=-0.18, SE=0.07, *t*(236.12)=-2.47, *p*=.014, 95% CI[−0.34, −0.03]), indicating that between-individual differences in average maturity, but not changes in pubertal stage between timepoints, moderated the association between testosterone and cortisol. Supporting our hypothesis, simple effects analyses at +/-1SD of the mean of average pubertal stage indicated that testosterone and cortisol were more strongly positively associated among adolescents in earlier puberty (β=0.60, SE=0.09, *t*(237.08)=6.38, *p*<.001, 95% CI[0.67, 0.87]) than they were among adolescents in later puberty (β=0.34, SE=0.08, *t*(230.61)=4.09, *p*<.001, 95% CI[0.18, 0.51]). Results of formal model fitting indicated that the main effect of sex and Δ pubertal stage improved model fit. When controlling for the main effect of sex, the interaction between testosterone and average pubertal stage was somewhat reduced in effect size and was no longer statistically significant (β=-0.14, SE=0.07, *t*(232.89)=-1.91, *p*=.058, 95% CI[−0.27, 0.01]).

In a separate model conducted within boys only, the interaction between testosterone and average pubertal stage was larger in effect size than it was in the full model including both boys and girls (β=-0.27, SE=0.10, 95% CI[−0.48, −0.06]). In contrast, in a separate model conducted within girls only, the interaction between testosterone and average pubertal stage approached zero (β=0.04 SE=0.11, 95% CI[−0.19, 0.29]). The simple associations between testosterone and cortisol at mean Tanner stages 1-2 and 3-5 for each sex are presented in Figure 2, Panel B.

### 3.3 Objective 2: Examine coupling of developmental increases in sex hormones with changes in cortisol

#### 3.3.1 DHEA

Supporting our hypothesis, larger within-individual increases in DHEA from T1 to T2 (Δ DHEA) were associated with larger increases in cortisol across this period (β=0.31, SE=0.05, *t*(127.38)=6.13, *p*<.001, 95% CI[0.21, 0.40]), above and beyond the significant positive effect of between-individual differences in average level of DHEA on cortisol (β=0.37, SE=0.07, *t*(148.65)=5.56, *p*<.001, 95% CI[0.24, 0.52]). Results were similar in models conducted within boys and girls separately.

#### 3.3.2 Testosterone

Contrary to our hypothesis, results were similar with respect with testosterone. Larger within-individual increases in testosterone from T1 to T2 (Δ testosterone) were also associated with larger increases in cortisol (β=0.31, SE=0.05, *t*(126.04)=6.43, *p*<.001, 95% CI[0.22, 0.41]), above and beyond the significant positive effect of between-individual differences in average level of testosterone (β=0.28, SE=0.07, *t*(136.46)=3.97, *p*<.001, 95% CI[0.14, 0.41]). Results were similar in models conducted within boys and girls separately.

### 3.4 Objective 3: Examine the effect of the severity of threat-related ELS on the coupling of developmental increases in sex hormones with changes in cortisol

#### 3.4.1 DHEA

The severity of threat-related ELS interacted with Δ DHEA from T1 to T2 to explain cortisol (β=-0.14, SE=0.05, *t*(124.29)=-2.51, *p*=.014, 95% CI[−0.25, −0.03]), but not with average levels of DHEA across timepoints (β=0.06, SE=0.06, *t*(131.41)=0.97, *p*=.332, 95% CI[−0.07, 0.18]). Contrary to our hypothesis, simple effects analyses at +/-1SD of the mean of threat-related ELS indicated that within-individual increases in DHEA were more strongly associated with increases in cortisol among adolescents who had been exposed to lower levels of threat-related ELS (β=0.46, SE=0.08, *t*(124.47)=5.94, *p*<.001, 95% CI[0.22, 0.43]) than they were among adolescents exposed to higher levels of threat-related ELS (β=0.19, SE=0.07, separately, the estimate for the interaction between Δ DHEA and threat-related ELS remained similar in girls (β=-0.18, SE=0.08, 95% CI[−0.33, −0.03]) but was considerably smaller in boys (β=-0.07, SE=0.07, 95% CI[−0.22, 0.08]). The severity of threat-related stress occurring between T1 and T2 did not interact with Δ DHEA or average levels of DHEA across timepoints to explain cortisol, suggesting a unique effect of stress occurring earlier in life. Model-estimated slopes for the sex-specific effects of within-individual changes in DHEA on changes in cortisol at higher and lower levels of threat-related ELS are presented in Figure 3, Panel A.

**Figure 3.**
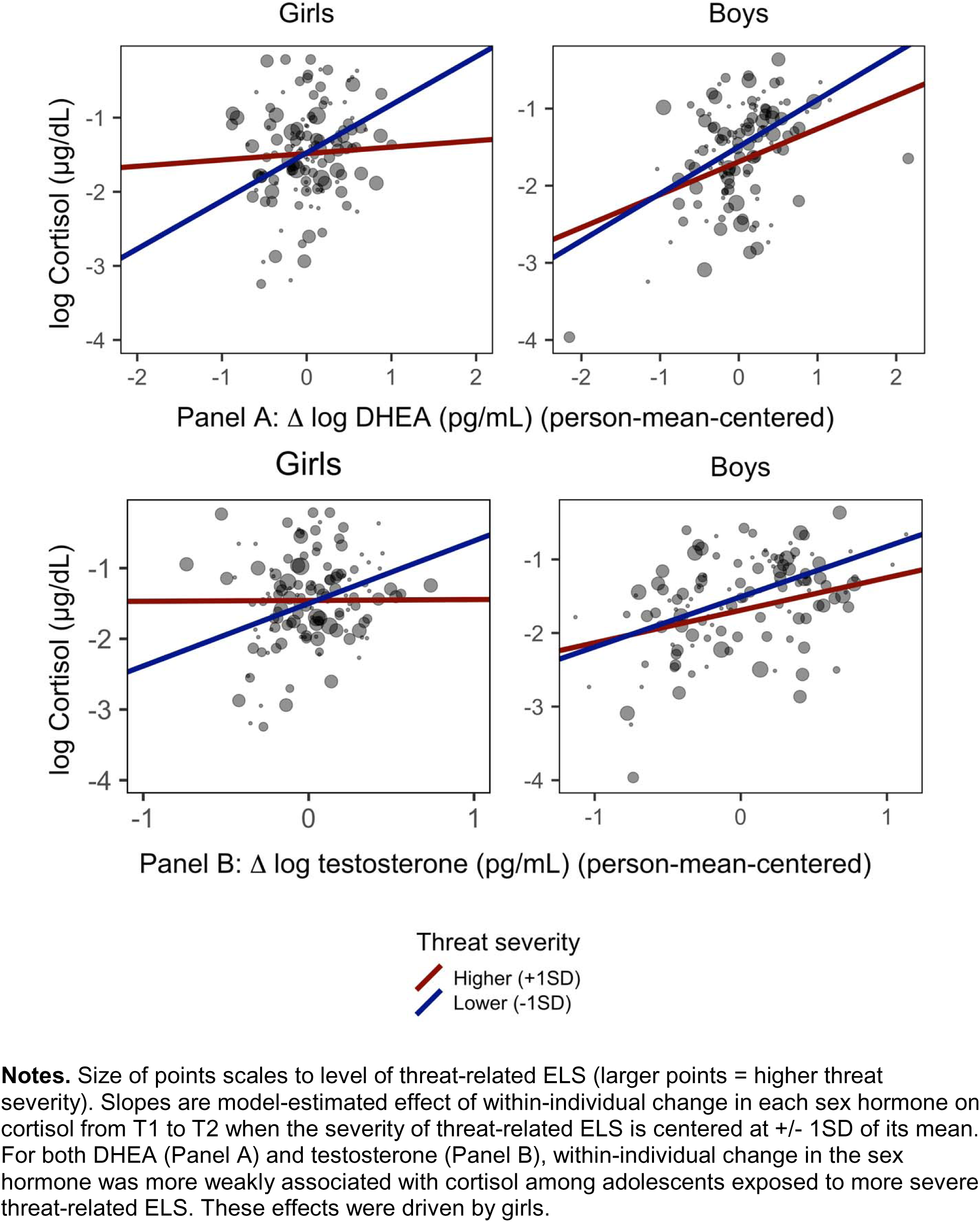
Association between changes in DHEA and testosterone and change in cortisol from T1 to T2 depends on the severity of threat-related ELS.

#### 3.4.2 Testosterone

Results were highly similar with respect to testosterone. The severity of threat-related ELS interacted with Δ testosterone from T1 to T2 (β=-0.12, SE=0.05, *t*(124.47)=-2.42, *p*=.017, 95% CI[−0.21, −0.02]) to explain cortisol, but not with average levels of testosterone across timepoints (β=-0.01, SE=0.06, *t*(131.43)=-0.20, *p*=.845, 95% CI[−0.15, 0.10]). Supporting our hypothesis, simple effects analyses at +/-1SD of the mean of the severity of threat-related ELS indicated that within-individual increases in testosterone were more strongly associated with increases in cortisol among adolescents who had been exposed to lower levels of threat-related ELS (β=0.42, SE=0.07, *t*(122.75)=6.42, *p*<.001, 95% CI[0.22, 0.40]) than they were among adolescents exposed to higher levels of threat-related ELS (β=0.19, SE=0.07, *t*(126.35)=2.72, *p*=.007, 95% CI[0.06, 0.33]). In models conducted within boys and girls separately, the estimate for the interaction of Δ testosterone with threat-related ELS remained similar in girls (β=-0.18, SE=0.08, 95% CI[−0.31, −0.02]) but was smaller in boys (β=-0.09, SE=0.06, 95% CI[−0.20, 0.03]). Once again, the severity of threat-related stress occurring between T1 and T2 did not interact with Δ testosterone or average levels of testosterone across timepoints to explain cortisol. Model-estimated slopes for the sex-specific effects of within-individual changes in testosterone on changes in cortisol at higher and lower levels of threat-related ELS are presented in Figure 3, Panel B.

## 4. Discussion

The goal of this study was to advance our understanding of the development of the HPA-axis across the transition through puberty. Using a two-timepoint longitudinal design, we first investigated the effects of between-individual levels and within-individual increases in pubertal stage on the association among waking levels of sex hormones and cortisol. Second, we examined whether developmental increases in waking levels of sex hormones are coupled with changes in waking levels of cortisol during puberty. Third, given prior evidence implicating both exposure to ELS and cortisol dysregulation in the development of mental and physical health difficulties (Koss and Gunnar, 2018), we examined whether the severity of exposure to threat-related ELS moderated the coupling of developmental increases in sex hormones with changes cortisol during puberty. Importantly, although we focused on *waking* levels of cortisol and sex hormones at two timepoints in development, several of our findings are consistent with those of previous studies examining the coupling of cortisol and sex hormones diurnally and in response to stress (Marceau et al., 2015b, 2014; Ruttle et al., 2015).

Findings from this study help to elucidate basic developmental processes concerning changes in and relations among cortisol, testosterone, and DHEA during puberty. In contrast to previous studies, we recruited and matched boys and girls based on early pubertal stage rather than on age. This design ensured that our findings were not confounded by sex differences in pubertal timing (Negriff and Susman, 2011) and increased our ability to examine the dynamics of cortisol and sex hormones that are specific to the pubertal transition. Specifically, because all boys and girls were in early puberty (self-reported Tanner stages 1-3) at the first timepoint, they advanced in puberty to varying degrees between the first and second timepoints. We found that the positive association between waking levels of cortisol and DHEA was stronger among adolescents who evidenced greater increases in pubertal stage across timepoints than it was among adolescents who evidenced smaller increases; between-individual differences in average level of pubertal stage did not moderate the association between levels of cortisol and DHEA. In contrast, levels of cortisol were more weakly positively associated with levels of testosterone in adolescents who were, on average, in later puberty compared to adolescents in earlier puberty; within-individual differences in the degree of change in pubertal stage across timepoints had no effect on the association between levels of cortisol and testosterone. These effects were driven by boys.

Most previous investigations of the associations of cortisol with sex hormones during adolescence have cross-sectionally compared adolescents at different ages rather than longitudinally examining changes in hormonal dynamics across the pubertal transition. Certainly, in these investigations older age was likely partially colinear with higher pubertal stage. Marceau et al. (2014) found that the cross-sectional positive association between age and cortisol–DHEA coupling in response to stress was driven by boys. In a longitudinal study, Ruttle et al. (2013) found that positive waking cortisol–DHEA coupling became stronger between ages 11 and 15 years. Our findings extend previous research by identifying within-individual increases in pubertal stage as a potential mechanism underlying strenthening of the positive association between levels of cortisol and DHEA during adolescence. Consistent with the findings of the current study, the association between testosterone and cortisol shows the opposite developmental pattern: investigators have found that cortisol–testosterone coupling is *weaker* (Marceau et al., 2014) or even negative (Ruttle et al., 2015) in older compared to younger adolescents. In the current study, many adolescents had not matured to the final stage of puberty by the second timepoint. A switch to negative coupling may occur later in pubertal development.

Positive associations among cortisol and sex hormones is likely due to the integrated synthesis and metabolism of cortisol and sex hormones by the HPA and HPG axes (Marceau et al., 2015a). Increasing strength of the positive association between cortisol and DHEA with advancing pubertal stage in boys suggests that puberty is a period of HPA-axis plasticity when recalibration occurs. To date, the mechanisms underlying pubertal plasticity of the HPA axis have not been elucidated. We found that when controlling for the significant positive effects of between-individual differences in levels of DHEA and testosterone on cortisol, within-individual increases in DHEA and testosterone across timepoints were positively coupled with increases in cortisol across this period. Thus, typical developmental increases in the hormones that are responsible for sexual maturation may contribute to the plasticity of cortisol regulation during puberty.

The few studies that have examined the effects of life stress on the associations of cortisol with sex hormones during adolescence have reported equivocal findings (Dismukes et al., 2015a, 2015b; Ruttle et al., 2015; Simmons et al., 2015). In the sole previous study examining the associations of cortisol with sex hormones across development, adolescents with low scores on a composite measure of threat- and deprivation-related ELS exhibited increases in the strength of waking cortisol–DHEA coupling across early adolescence; in contrast, adolescents with high scores on this measure exhibited initial increases in the strength of coupling, which then plateaued (Ruttle et al., 2015). In a cross-sectional study of diurnal coupling, Marceau et al. (2015b) found that deviations in DHEA and testosterone from the diurnal slope were positively associated with deviations in cortisol, suggesting that coupling between cortisol and sex hormones may be due to similar responses to events in the daily environment. Contrary to our hypothesis and the formulation that environmental threat would accelerate development and result in stronger positive coupling of increases in DHEA and cortisol across timepoints, we found that higher severity of exposure to threat-related ELS was associated with *attenuated* coupling of increases in DHEA and in cortisol. It is possible, therefore, that typical environmental input (e.g., minor daily stressors) results in concordant deviations in cortisol and DHEA, whereas environmental input that is characterized by severe threat reduces developmentally-typical concordance. Consistent with our hypothesis, we found that greater severity of exposure to threat-related ELS was associated with weaker coupling of changes in testosterone and cortisol; however, the interpretation that this pattern reflects “stress acceleration” is questionable given that the effect of threat-related ELS cortisol–DHEA coupling was opposite to the hypothesized direction.

Finally, we found no effect of the severity of threat-related stress occurring *during puberty* on coupling of developmental increases in sex hormones and cortisol, suggesting a unique effect of stress occurring earlier in life. Previous studies examining pubertal recalibration of cortisol regulation have focused on previously institutionalized children who experienced dramatic shifts in the quality of the environment from early to late childhood and subsequently showed normalization of cortisol regulation during puberty (DePasquale et al., 2019; Gunnar et al., 2019; Quevedo et al., 2012). Among children raised in their families of origin, however, the quality of the environment is likely to remain consistent across childhood and adolescence (Dunn et al., 2011); indeed, in our sample, the severity of threat-related ELS was correlated with the severity of threat-related stress occurring during the transition through puberty. Thus, for most children, puberty is likely to be a period during which the effects of exposure to severe ELS on HPA-axis functioning emerge or intensify, rather than remediate.

We should note four primary limitations of this study. First, although this study was longitudinal, we assessed adolescents at only two timepoints; therefore, we were unable to model *trajectories* of the associations of cortisol and sex hormones during puberty, which may not be linear. While our study design allowed us to examine whether changes in sex hormones predicted changes in cortisol from early to later puberty above and beyond levels of these hormones in early puberty, we cannot draw causal conclusions concerning whether changes in sex hormones *drive* changes in cortisol. Instead, changes in cortisol during puberty may drive changes in sex hormones, although this explanation is less likely given that our design focused on a developmental period in which sex hormones increased substantially. Second, in addition to advancing in pubertal stage, boys and girls chronologically aged from the first to second timepoints, and it is possible that maturational processes separate from pubertal development partially contributed to the observed changes in hormonal dynamics. It is challenging to disentangle the influence of these processes from pubertal maturation. Third, in order to recruit adolescents who were in early puberty at the baseline assessment, we excluded girls who had started menses. Given that ELS has been associated with earlier menarche (e.g., Chisholm, Quinlivan, Petersen, & Coall, 2005) we may have selected a subset of stress-exposed girls who were less vulnerable to the effect of ELS on the timing of menarche.

A third limitation of this study is that we assessed waking levels of hormones on a single day at each timepoint. Importantly, single measures of hormones may not be reliable measures of typical functioning given that averaging multiple measures of the same construct generally yields higher reliability (Kraemer et al., 2001). We should note, however, that concerns regarding the reliability of the hormone values in the current study are mitigated by the fact that the hormone values showed moderate stability and agreement across the two timepoints despite anticipated developmental change, that sex hormones increased as expected, and that our findings were largely consistent with those reported in previous studies. Nonetheless, we could not quantify the reliability of hormone values within each timepoint. Lower reliability of the hormone values due to relying on single samples would compromise the construct validity of our measures. We measured waking hormone levels in order to minimize the influence of unmeasured variables (e.g., minor daily stress); nevertheless, we were unable to examine changes in the dynamics of the associations of cortisol with sex hormones across the day from early to later puberty. Further, although adolescents and their parents were provided with instructions to collect salivary hormone samples immediately upon awakening, we had no way of verifying participant compliance with the instructions. Given that cortisol typically rises following waking (i.e., the cortisol awakening response), unmeasured systematic differences in participant compliance could have biased some of our results.

The implications of HPA-axis dysregulation for mental and physical health and the effects of life stress on HPA-axis regulation have been studied for decades (Koss and Gunnar, 2018). The vast majority of previous studies, however, have focused on cortisol production in isolation, and have not examined the associations of cortisol with other related hormones. This gap in the literature has hampered our knowledge of the development of the HPA axis, and, ultimately, of the relation of HPA-axis functioning with long-term mental and physical health outcomes. Findings of the current study indicate that the association of sex hormones with cortisol changes as a function of pubertal maturation and that that exposure to threat-related ELS influences coupling of pubertal increases in sex hormones with changes in cortisol. These findings elucidate mechanisms underlying the pubertal recalibration of the HPA axis.

## Supporting information

Supplementary Text, Tables, and Analyses

## Disclosures and Acknowledgements

The authors report no conflicts of interest.

This research was supported by the National Institutes of Health (F32-MH114317 to NLC; R37-MH101495 to IHG) and the National Science Foundation Graduate Fellowship Awards (to NLC and LSK). We thank Alexandria Price, Holly Pham, Isabella Lazzareschi, Cat Camacho, Monica Ellwood-Lowe, Sophie Schouboe, Maddie Pollak, Morgan Popolizio, Anna Chichocki, Lucinda Sisk, Rachel Weisenburger, and Michelle Sanabria for their assistance in scheduling and running participants. We also thank the participants and their families for their contributions to this project.

## References

Adam, E.K., Quinn, M.E., Tavernier, R., McQuillan, M.T., Dahlke, K.A., Gilbert, K.E., 2017. Diurnal cortisol slopes and mental and physical health putcomes:A systematic review and meta-analysis. Psychoneuroendocrinology 83, 25–41. https://doi.org/10.1016/j.psyneuen.2017.05.018

Bates, D., Machler, M., Bolker, B., Walker, S., 2014. Fitting linear mixed-effects models using lme4. J. Stat. Softw. 1–51.

Bernard, K., Frost, A., Bennett, C.B., Lindhiem, O., 2017. Maltreatment and diurnal cortisol regulation: A meta-analysis. Psychoneuroendocrinology 78, 57–67. https://doi.org/10.1016/j.psyneuen.2017.01.005

Black, S.R., Lerner, M.D., Shirtcliff, E.A., Klein, D.N., 2018. Patterns of neuroendocrine coupling in 9-year-old children: Effects of sex, body-mass index, and life stress. Biol. Psychol. 132, 252–259. https://doi.org/10.1016/j.biopsycho.2017.11.004

Bordini, B., Rosenfield, R.L., 2011. Normal pubertal development: Part I: The endocrine basis of puberty. Pediatr. Rev. 32, 223–229.

Bunea, I.M., Szentágotai-Tă, A., Miu, A.C., 2017. Early-life adversity and cortisol response to social stress: A meta-analysis. Transl. Psychiatry 7, 1274. https://doi.org/10.1038/s41398-017-0032-3

Chambers, J.M., 1992. Linear models, in: Chambers, J.M., Hastie, T.J. (Eds.), Statistical Models in S. Wadsworth & Brooks/Cole.

Chisholm, J.S., Quinlivan, J.A., Petersen, R.W., Coall, D.A., 2005. Early stress predicts age at menarche and first birth, adult attachment, and expected lifespan. Hum. Nat. 16, 233–265. https://doi.org/10.1007/s12110-005-1009-0

Colich, N.L., Kircanski, K., Foland-Ross, L.C., Gotlib, I.H., 2015. HPA-axis reactivity interacts with stage of pubertal development to predict the onset of depression. Psychoneuroendocrinology 55, 94–101. https://doi.org/10.1016/j.psyneuen.2015.02.004

Colich, N.L., Rosen, M.L., Williams, E.S., McLaughlin, K.A., 2019. Biological aging in childhood and adolescence following experiences of threat and deprivation: A systematic review and meta-analysis. bioRxiv 1–85.

Curran, P.J., Bauer, D.J., 2011. The Disaggregation of Within-Person and Between-Person Effects in Longitudinal Models of Change. Annu. Rev. Psychol. 62, 583–619. https://doi.org/10.1146/annurev.psych.093008.100356.The

Del Giudice, M., Ellis, B.J., Shirtcliff, E.A., 2011. The Adaptive Calibration Model of stress responsivity. Neurosci. Biobehav. Rev. 35, 1562–1592. https://doi.org/10.1016/j.neubiorev.2010.11.007

DePasquale, C.E., Donzella, B., Gunnar, M.R., 2019. Pubertal recalibration of cortisol reactivity following early life stress: a cross-sectional analysis. J. Child Psychol. Psychiatry 60, 568– 575. https://doi.org/10.1111/jcpp.12992

Dismukes, A.R., Johnson, M.M., Vitacco, M.J., Iturri, F., Shirtcliff, E.A., 2015a. Coupling of the HPA and HPG axes in the context of early life adversity in incarcerated male adolescents. Dev. Psychobiol. 57, 705–718. https://doi.org/10.1002/dev.21231

Dismukes, A.R., Shirtcliff, E.A., Hanson, J.L., Pollak, S.D., 2015b. Context influences the interplay of endocrine axes across the day. Dev. Psychobiol. 57, 731–741. https://doi.org/10.1002/dev.21331

Dorn, L.D., Biro, F.M., 2011. Puberty and its measurement: A decade in review. J. Res. Adolesc. 21, 180–195. https://doi.org/10.1111/j.1532-7795.2010.00722.x

Dunn, V.J., Abbott, R.A., Croudace, T.J., Wilkinson, P., Jones, P.B., Herbert, J., Goodyer, I.M., 2011. Profiles of family-focused adverse experiences through childhood and early adolescence: The ROOTS project a community investigation of adolescent mental health. BMC Psychiatry 11, 109. https://doi.org/10.1186/1471-244X-11-109

Essex, M.J., Shirtcliff, E.A., Burk, L.R., Ruttle, P.L., Klein, M.H., Slattery, M.J., Kalin, N.H., Armstrong, J.M., 2011. Influence of early life stress on later hypothalamic-pituitary-adrenal axis functioning and its covariation with mental health symptoms: a study of the allostatic process from childhood into adolescence. Dev. Psychopathol. 23, 1039–58. https://doi.org/10.1017/S0954579411000484

Gunnar, M., Quevedo, K., 2007. The neurobiology of stress and development. Annu. Rev. Psychol. 58, 145–173. https://doi.org/10.1146/annurev.psych.58.110405.085605

Gunnar, M.R., DePasquale, C.E., Reid, B.M., Donzella, B., 2019. Pubertal stress recalibration reverses the effects of early life stress in postinstitutionalized children. Proc. Natl. Acad. Sci. U. S. A. 1–5. https://doi.org/10.1073/pnas.1909699116

Hucklebridge, F., Hussain, T., Evans, P., Clow, A., 2005. The diurnal patterns of the adrenal steroids cortisol and dehydroepiandrosterone (DHEA) in relation to awakening. Psychoneuroendocrinology 30, 51–57. https://doi.org/10.1016/j.psyneuen.2004.04.007

Humphreys, K.L., Kircanski, K., Colich, N.L., Gotlib, I.H., 2016. Attentional avoidance of fearful facial expressions following early life stress is associated with impaired social functioning. J. Child Psychol. Psychiatry 57, 1174–1182. https://doi.org/10.1111/jcpp.12607

Johnson, M.M., Dismukes, A.R., Vitacco, M.J., Breiman, C., Fleury, D., Shirtcliff, E.A., 2014. Psychopathy’s influence on the coupling between hypothalamic-pituitary-adrenal and - gonadal axes among incarcerated adolescents. Dev. Psychobiol. 56, 448–458. https://doi.org/10.1002/dev.21111

Kamin, H.S., Kertes, D.A., 2016. Cortisol and DHEA in development and psychopathology. Horm. Behav. 89, 69–85. https://doi.org/10.1016/j.yhbeh.2016.11.018

King, L.S., Colich, N.L., LeMoult, J., Humphreys, K.L., Ordaz, S.J., Price, A.N., Gotlib, I.H., 2017. The impact of the severity of early life stress on diurnal cortisol: The role of puberty. Psychoneuroendocrinology 77, 68–74. https://doi.org/10.1016/j.psyneuen.2016.11.024

King, L.S., Humphreys, K.L., Camacho, M.C., Gotlib, I.H., 2019. A person-centered approach to the assessment of early life stress: Associations with the volume of stress-sensitive brain regions in early adolescence. Dev. Psychopathol. 31, 643–655. https://doi.org/10.1017/S0954579418000184

Koss, K.J., Gunnar, M.R., 2018. Early adversity, the hypothalamic-pituitary-adrenocortical axis, and child psychopathology. J. Child Psychol. Psychiatry 59, 327–346. https://doi.org/10.1111/jcpp.12784

Kraemer, H.C., Ph, D., Stice, E., Kazdin, A., Offord, D., Kupfer, D., 2001. How do risk factors work together? Mediators, moderators, and independent, overlapping, and proxy risk factors. Psychiatry Interpers. Biol. Process. 158, 848–856. https://doi.org/10.1176/appi.ajp.158.6.848

Kuznetsova, A., Brockhoff, P.B., Christensen, R.H.B., 2016. Tests in linear mixed effects models. J. Stastical Softw. 82, 1–26. https://doi.org/10.18637/jss.v082.i13

Lambert, H.K., King, K.M., Monahan, K.C., McLaughlin, K.A., 2017. Differential associations of threat and deprivation with emotion regulation and cognitive control in adolescence. Dev. Psychopathol. 29, 929–940. https://doi.org/10.1017/S0954579416000584

Maninger, N., Wolkowitz, O.M., Reus, V.I., Epel, E.S., Mellon, S.H., 2009. Neurobiological and neuropsychiatric effects of dehydroepiandrosterone (DHEA) and DHEA sulfate (DHEAS). Front. Neuroendocrinol. 30, 65–91. https://doi.org/10.1016/j.yfrne.2008.11.002

Marceau, K., Ruttle, P.L., Shirtcliff, E.A., Essex, M.J., Susman, E.J., 2015a. Developmental and contextual considerations for adrenal and gonadal hormone functioning during adolescence: Implications for adolescent mental health. Dev. Psychobiol. 57, 742–768. https://doi.org/10.1002/dev.21214

Marceau, K., Ruttle, P.L., Shirtcliff, E.A., Hastings, P.D., Klimes-Dougan, B., Zahn-Waxler, C., 2015b. Within-person coupling of changes in cortisol, testosterone, and DHEA across the day in adolescents. Dev. Psychobiol. 57, 654–669. https://doi.org/10.1002/dev.21173

Marceau, K., Shirtcliff, E.A., Hastings, P.D., Klimes-Dougan, B., Zahn-Waxler, C., Dorn, L.D., Susman, E.J., 2014. Within-adolescent coupled changes in cortisol with DHEA and testosterone in response to three stressors during adolescence. Psychoneuroendocrinology 41, 33–45. https://doi.org/10.1016/j.psyneuen.2013.12.002

Marshall, W.A., Tanner, J., 1968. Growth and physiological development during adolescence. Annu. Rev. Med. 19, 283–300.

Mastorakos, G., Pavlatou, M.G., Mizamtsidi, M., 2006. The hypothalamic-pituitary-adrenal and the hypothalamic-pituitary-gonadal axes interplay. Pediatr. Endocrinol. Rev. 3 Suppl 1, 172–81.

Matchock, R.L., Dorn, L.D., Susman, E.J., 2007. Diurnal and seasonal cortisol, testosterone, and DHEA rhythms in boys and girls during puberty. Chronobiol. Int. 24, 969–990. https://doi.org/10.1080/07420520701649471

McEwen, B.S., 2019. What is the confusion with cortisol? Chronic Stress 3, 1–3. https://doi.org/10.1177/2470547019833647

Morris, N.M., Udry, J.R., 1980. Validation of a self-administered instrument to assess stage of adolescent development. J. Youth Adolesc. 9, 271–280. https://doi.org/10.1007/BF02088471

Negriff, S., Susman, E.J., 2011. Pubertal timing, depression, and externalizing problems: A framework, review, and examination of gender differences. J. Res. Adolesc. 21, 717–746. https://doi.org/10.1111/j.1532-7795.2010.00708.x

Quevedo, K., Johnson, A., Loman, M., Lafavor, T., Gunnar, M., 2012. The confluence of adverse early experience and puberty on the cortisol awakening response. Int. J. Behav. Dev. 36, 19–28. https://doi.org/10.1177/0165025411406860.The

R Core Team, 2018. R: A language and environment for statistical computing.

Romeo, R.D., 2013. The teenage brain: The stress response and the adolescent brain. Curr. Dir. Psychol. Sci. 22, 140–145. https://doi.org/10.1177/0963721413475445

Ruttle, P.L., Shirtcliff, E.A., Armstrong, J.M., Klein, M.H., Essex, M.J., 2015. Neuroendocrine coupling across adolescence and the longitudinal influence of early life stress. Dev. Psychobiol. 57, 688–704. https://doi.org/10.1002/dev.21138

Sapolsky, R.M., 1997. McEwen-induced modulation of endocrine history: A partial review. Stress 2, 1–11. https://doi.org/10.3109/10253899709014733

Sheridan, M.A., McLaughlin, K.A., 2014. Dimensions of early experience and neural development: Deprivation and threat. Trends Cogn. Sci. 18, 580–585. https://doi.org/10.1016/j.tics.2014.09.001

Sheridan, M.A., Peverill, M., Finn, A.S., McLaughlin, K.A., 2017. Dimensions of childhood adversity have distinct associations with neural systems underlying executive functioning. Dev. Psychopathol. 29, 1777–1794. https://doi.org/10.1017/S0954579417001390

Simmons, J.G., Byrne, M.B., Schwartz, O.S., Whittle, S.L., Sheeber, L., Kaess, M., Youssef, G.J., Allen, N.B., 2015. Dual-axis hormonal covariation in adolescence and the moderating influence of prior trauma and aversive maternal parenting. Dev. Psychobiol. 57, 670–687. https://doi.org/10.1002/dev.21275

Woltman, H., Feldstain, A., Mackay, J.C., Rocchi, M., 2012. An introduction to hierarchical linear modeling. Tutor. Quant. Methods Psychol. 8, 52–69.

