## Supplementary Text, Tables, and Analyses for "Associations of waking cortisol with DHEA and testosterone across the pubertal transition: Effects of threat-related early life stress"

**Method**

**Procedure**

The T2 assessment occurred an average of 2 years following the T1 assessment (mean[SD]=1.96[0.31] years; range: 1.18-2.88). The timing of the T2 assessment varied for a variety of reasons related to the challenges of collecting longitudinal data in a large adolescent sample. The target date for the T2 assessment was 18 months following the T1 assessment. However, many participants could not be reached or scheduled in time and were thus assessed >18 months. Further, because the larger study involved an MRI session, participants who received metal braces between T1 and T2 could not participate until these braces had been removed. The participants who were assessed <18 months after T1 were those who we assessed early because they notified us that they would soon be receiving braces. The distribution of the interval (in years) between T1 and T2 is presented in Figure S1.

**Figure S1. Distribution of interval (in years) between T1 and T2**

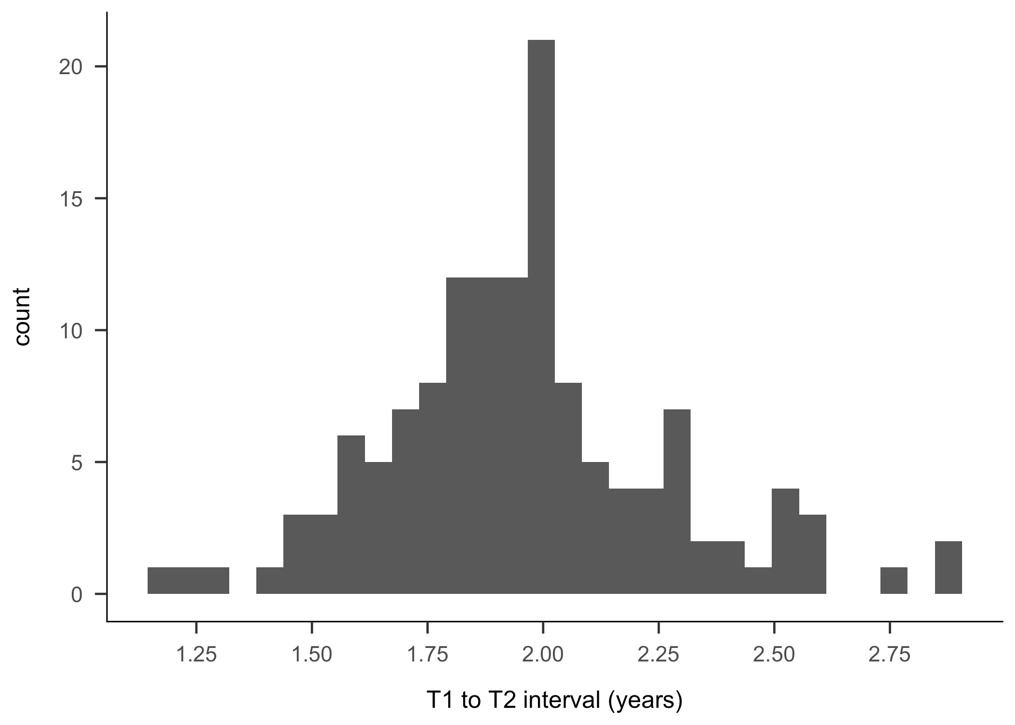

### Measures

**Pubertal stage.** Distributions of Tanner scores at each time-point and change in Tanner scores from T1 to T2 are presented in Figures S2 and S3.

**Figure S2. Distribution of self-reported Tanner staging scores at each time-point**

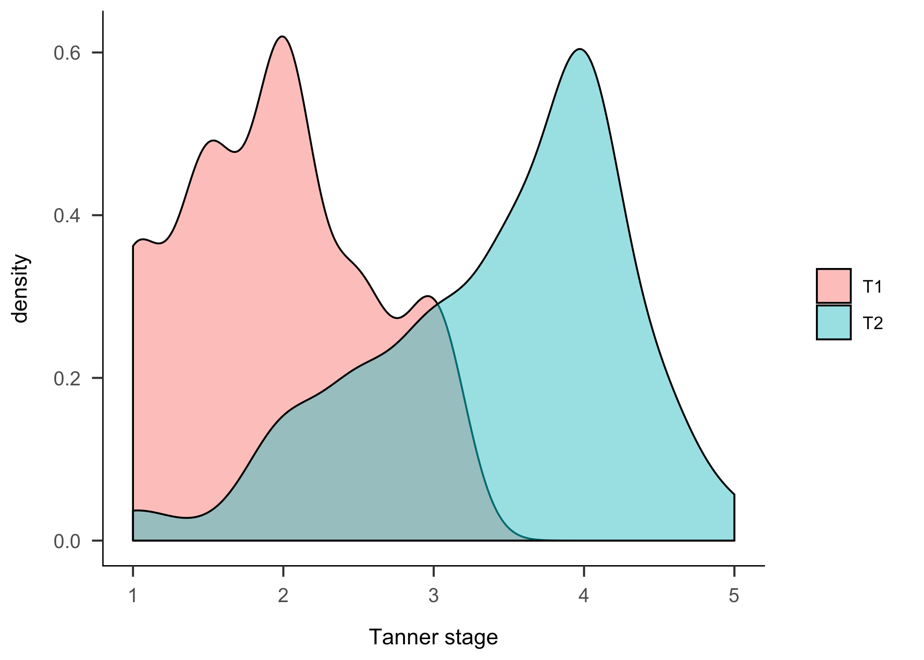

**Figure S3. Distribution of change in self-reported Tanner staging scores from T1 to T2**

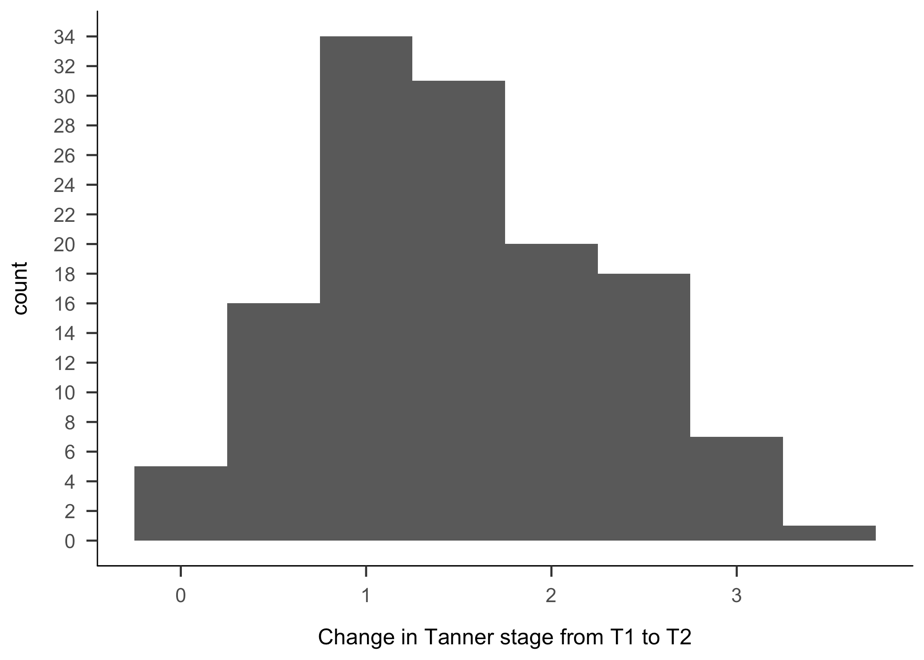

**Table S1.** Endorsement of threat-related stressors occurring between T1 and T2

| Type of ELS | N  endorsed | %  endorsed | Mean (SD) Severity |
| --- | --- | --- | --- |
| Family verbal conflict | 32 | 22 | 1.88(0.68) |
| Bullying | 23 | 16 | 1.35(0.57) |
| Community violence | 15 | 10 | 0.63(0.61) |
| Community instability | 5 | 3 | 0.80(0.45) |
| Domestic violence | 3 | 2 | 2.34(1.26) |
| Community verbal conflict | 10 | 7 | 0.50(0.34) |
| Emotional abuse | 9 | 6 | 2.39(0.60) |
| Physical abuse | 0 | 0 | N/A |
| Mugging or robbery | 5 | 3 | 1.00(0.61) |
| War or terrorism | 3 | 2 | 0.50(0.00) |
| Sexual abuse | 0 | 0 | N/A |
| Threats of domestic violence | 1 | <1 | 1.50(N/A) |
| Kidnapping | 0 | 0 | N/A |
| Threats of physical abuse | 1 | <1 | 1.50(N/A) |
| Witness sexual abuse | 0 | 0 | N/A |

**Notes.** Stressors coded as **“**community instability” included community-level threats (e.g., bomb/active shooter threats at school, hearing gun shots in neighborhood). Stressors coded as “war or terrorism” included witnessing events live on television.

**Waking cortisol, DHEA, and testosterone.**

To increase compliance, parents were also provided with written and oral instructions for sample collection. The high sensitivity enzyme immunoassays were as follows: Cat No. 1-3002 for cortisol; Cat. No. 1-1202 for DHEA; Cat. No. 1-2402 for testosterone. The assay for cortisol used 25 μl of saliva per determination, had a lower limit sensitivity of .0007 µg/dL, and a standard curve range from .012-3.0 µg/dL. The assay for DHEA used 50 μl of saliva per determination, had a lower limit of sensitivity of 5 pg/mL, and a standard curve range from 10.2-1000 pg/mL. The assay for testosterone used 25 μl of saliva per determination, had a lower limit of sensitivity of 1 pg/mL, and a standard curve range from 6.1-600 pg/mL.

Of the 200 adolescents who met eligibility criteria at T1, 26 did not provide a saliva sample at T1 and 3 provided samples that were of insufficient quantity for measurement of any hormone. Of the remaining 171 adolescents who provided usable cortisol, DHEA, *or* testosterone values at T1, 151 provided cortisol (19 insufficient quantity, 1 below lower limit of assay kit), 170 provided DHEA (1 below lower limit of assay kit), and 168 provided testosterone (3 insufficient quantity). Of the 200 adolescents who met eligibility criteria at T1, 32 were lost to follow-up, 13 withdrew from the study, 4 did not provide a saliva sample at T2, 2 opted not to participate at T2, and 2 provided samples that were of insufficient quantity for measurement of hormone. Of the remaining 147 adolescents who provided usable cortisol, DHEA, *or* testosterone values at T2, 140 provided cortisol (7 insufficient quantity), 144 provided DHEA (3 insufficient quantity), and all 147 provided testosterone.

**Results**

**Main effects of pubertal stage on change in cortisol, DHEA, and testosterone**

Results of multi-level models regressing the hormone values onto both within-person longitudinal change in pubertal stage and mean pubertal stage across time-points are presented in Table S2. For both DHEA and testosterone *both* larger within-person increases in pubertal stage between T1 and T2 and higher mean pubertal stage across time-points were associated with higher hormones. In contrast, for cortisol, only larger increases in pubertal stage were associated with higher cortisol.

**Table S2.** Effects of within-person longitudinal change in pubertal stage and between-person average pubertal stage on cortisol, DHEA, and testosterone

| **Outcome** | **Predictor** | **β (SE)** | ***df*** | ***t*** | ***p*** | **95% CI** |
| --- | --- | --- | --- | --- | --- | --- |
| cortisol | Δ pubertal stage | 0.16 (0.06) | 122.92 | 2.80 | .006 | 0.05, 0.28 |
|  | mean pubertal stage | 0.07 (0.08) | 127.91 | 0.86 | .393 | -0.08, 0.23 |
| DHEA | Δ pubertal stage | 0.16 (0.04) | 130.67 | 3.72 | <.001 | 0.08, 0.24 |
|  | mean pubertal stage | 0.26 (0.08) | 129.17 | 3.51 | <.001 | 0.11, 0.43 |
| testosterone | Δ pubertal stage | 0.36 (0.04) | 131.34 | 8.50 | <.001 | 0.29, 0.45 |
|  | mean pubertal stage | 0.27 (0.07) | 129.79 | 3.89 | <.001 | 0.13, 0.41 |

**Notes.** Δ pubertal stage = person-mean-centered Tanner stage at T1 and T2; mean pubertal stage = mean Tanner stage across T1 and T2; SE = standard error; df = degrees of freedom; CI = bootstrapped confidence interval for β.

**Sex differences in cortisol, DHEA, and testosterone and in the effects of pubertal stage on change in these hormones**

Girls had significantly higher levels of cortisol at T1 than did boys (β=0.22, SE=0.08, *t*(279.59)=2.81, *p*=.005, 95% CI[0.07, 0.38]). Further, multi-level models indicated that sex interacted with within-person Δ pubertal stage to explain cortisol (β=0.28, SE=0.11, *t*(122.99)=2.49, *p*=.014, 95% CI[0.06, 0.49]), such that increases in pubertal stage between T1 and T2 were associated with increases in cortisol in boys (β=0.28, SE=0.07, *t*(117.87)=3.78, *p*<.001, 95% CI[0.14, 0.43]), but not in girls (β=<0.01, SE=0.08, *t*(127.29)=0.84, *p*=.968, 95% CI[-0.16, 0.16]). At T1, boys and girls did not differ significantly in levels of testosterone (β=0.09, SE=0.07, *t*(270.55)=1.39, *p*=.167, 95% CI[-0.04, 0.22]); however, at T2, boys had significantly higher levels of testosterone than did girls (β=0.59, SE=0.07, *t*(285.81)=8.53, *p*<.001, 95% CI[0.45, 0.72]). Sex interacted with within-person Δ pubertal stage (β=0.32, SE=0.06, *t*(130.88)=5.55, *p*<.001, 95% CI[0.20, 0.43]) to explain testosterone, such that increases in pubertal stage were associated with higher testosterone for both boys and girls; however, this association was stronger in boys (β=0.57, SE=0.05, *t*(132.61)=10.60, *p*<.001, 95% CI[0.47, 0.68]) than in girls (β=0.14, SE=0.06 *t*(129.31)=2.45, *p*=.016, 95% CI[0.03, 0.25]). Finally, greater within-person Δ pubertal stage was associated with larger increases in DHEA between T1 and T2 for both boys and girls (β=0.16, SE=0.04, *t*(130.67)=3.72, *p*<.001, 95% CI[0.08, 0.24]). There were no sex differences in levels of DHEA at either time-point nor in the within- and between-person effects of pubertal stage on change in DHEA across time-points. Sex differences in cortisol, DHEA, and testosterone at each time-point and as a function of Δ pubertal stage are presented in Figure 1.

**Effects of the severity of non-threat-related ELS on the coupling of developmental increases in sex hormones with changes in cortisol** In the primary analysis, we focused on threat-related ELS in order to increase the specificity of our hypotheses. The severity of non-threat-related ELS was quantified as the sum of the maximum severity rating for each for all of the events not included in the threat-related ELS calculation (i.e., the events that were not interpersonal in nature and/or did not meet the Sheridan and McLaughlin (2014) definition of threat). These events were: death of someone close, parental divorce/separation, disaster, animal attack, witnessing or experiencing an accident or illness, family financial problems, family legal problems, moving, mental illness of someone close, neglect, separation from family, and suicide completion or attempt of someone close. There were no effects of non-threat-related ELS on the association between within-individual increases in DHEA or between-person levels of DHEA and change in cortisol across time-points (*p*-values > .239). Similarly, there were no effects of non-threat-related ELS on the association between within-individual increases in testosterone or between-person levels of testosterone and change in cortisol across time-points (*p*-values > .323).

**References**

Sheridan, M. A., & McLaughlin, K. A. (2014). Dimensions of early experience and neural development: Deprivation and threat. *Trends in Cognitive Sciences*, *18*(11), 580–585. https://doi.org/10.1016/j.tics.2014.09.001
